# Average stride length and stride rate of Thoroughbreds and Quarter Horses during ‘Sprint’ and ‘Classic’ races

**DOI:** 10.1101/2021.07.27.453998

**Authors:** Fernando B. Vergara-Hernandez, Brian D. Nielsen, Cara I. Robison, Taylor A. Fabus, Jasmin L. Kompare, Rebecca A. LeCompte Lasic, Aimee C. Colbath

## Abstract

The main factors influencing speed in athletes are stride length (SL) and stride rate (SR). However, conflict remains whether SL or SR is the key determinant of higher speeds. Quarter Horses (QH) generally reach higher speeds in their races than do Thoroughbreds (TB). However, the influence of SL and SR on this greater speed is unclear. Therefore, the main objective of this study was to compare SL and SR in QH and TB raced in short (sprint) and long (classic) distances. We hypothesized that QH have a higher SR in comparison to TB, and SR decreases as distance increases. Two race distances were analyzed for each breed: QH races of 100.6 and 402.3 m, and TB races of 1,207.0 m and 2,011.7 m. Data from twenty horses were obtained, consisting of five horses from each race distance (10 QH and 10 TB). Five individuals watched recordings of each race three times counting the number of strides taken by each winning horse. The SR was calculated using the average number of strides over a given race duration, and SL was determined by calculating the total number of strides over the distance covered. Speed was calculated dividing the distance by the time of the winning horse. The PROC Mixed Procedure was used to identify statistical differences between breeds, and between distances within the same breed. Results showed that although the SL of the TB was longer in comparison with the QH (P<0.001), the average SR in QH was higher than in TB (2.88 vs 2.34 + 0.03 strides/s; P<0.001). Further, QH classic distance demonstrated a faster speed than TB at either distance (P<0.001). In conclusion, QH achieve a higher SR in comparison to TB (between 14-20% more than TB), confirming the importance of SR in achieving high racing speeds.

## Introduction

Whether it is a human sprinter, a marathoner, or a racehorse, speed is critical to performance. It is a common belief among sports enthusiasts that a greater stride length (SL) will result in a faster individual. While many have been focusing only on SL, velocity is dictated by SL and SR [1]. Daniels [2] identified that elite human athletes maximize their performance by reaching an average step rate of 180 steps/min (90 strides/min) or more in racing distances of 800 m or more. Some believe that SL and SR may have an inverse relationship [3]. In horses, it is currently unknown whether SL or SR is most influential in racing speed. Determining the importance of SL or SR on racing speed could have far reaching implications on horse selection.

Similarly, horse enthusiasts often believe that SL is the most important factor influencing speed. For instance, the outstanding racehorses Man O’ War, Secretariat, and Justify have SL reported to be 8.5, 7.6, and 7.5 m respectively [4]. These stride lengths are often quoted, leading to the potential misconception that speed in horses is most strongly influenced by SL. Thoroughbreds (TB) are the world’s most widely distributed racehorse and are well-known for their speed [5]. Triple crown winners have achieved peak speeds of 61.2 km/h [6]. However, this is not the fastest horse breed. Quarter Horses (QH), developed in the United States, are the fastest horse to run a quarter of a mile (403.3 m) [7], reaching top speeds of 87.5 to 92.6 km/h [7, 8]. QH and TB typically race different distances. For instance, QH most often run distances between 91-796 m, and TB usually run between 1,006-3,219 m (5-16 furlongs) [9].

Previous studies investigating the effect of SL and SR on racing speed have conflicting results. An early study showed that SL was responsible for the increase in speed, up to 8.3 m/s [10]. However, at higher speeds (11.7 m/s), increasing SR resulted in faster speeds [10]. Another study using three TB at six different speeds reported that, as the speed increases, both SL and SR increased nearly linearly. Yet, the fastest TB from the study had the shortest SL and highest SR at maximum speed indicating that, at higher speeds, SL may decrease as SR increases [11]. In another study using nine TB, it was observed that the SR increased linearly without signs of a plateau [12].

The objective of the current study was two-fold. The first objective was to evaluate the average SL and SR in both QH and TB breeds within their breed-specific races. The second objective was to determine if SL and SR are influenced by the distance raced. Therefore, it was hypothesized that QH have a greater average SR than TB, and that SR will decrease as running distance increases. Finally, it was hypothesized that SL would decrease as the SR increases.

## Material and Methods

To examine differences between breeds and between short and long race distances, “sprint” races and “classic” races were analyzed for both QH and TB from 2008 through 2012. The QH sprint race analyzed was the Texas Twister Stakes, 100.6 m (110 yards), held at Sam Houston Race Park (www.shrp.com), and the QH classic race was the Champion of Champions, 402.3 m (440 yards), held at Los Alamitos Race Course (www.losalamitos.com). For the TB races, the Breeders’ Cup Sprint, 1,207.0 m (6 furlongs), and Breeders’ Cup Classic, 2,011.7 m (10 furlongs), were analyzed. The Breeders’ Cup races were held at Santa Anita Park (www.santaanita.com) in 2008, 2009, and 2012, and at Churchill Downs (www.churchilldowns.com) in 2010 and 2011 [13]. This resulted in a total of 20 horses being analyzed (5 QH sprint, 5 QH classic, 5 TB sprint, 5 TB classic). Five individuals watched recordings of each race three times (in slow motion when necessary) and counted the strides of the winning horse. If, during periods of a race, the winning horse was not in camera view, the horse in the lead was chosen as a surrogate horse for which to count strides until the winning horse was back in the camera view. From the 15 viewings of each race (five individuals viewed each race three times), the average number of strides for the winning horse was calculated.

For the QH races, the stride count began when the starting gates opened, signifying the official start to the race. For TB races, the stride count began after the horses left the starting gates, had completed the “run-up” (passed the ‘flagman’ or ‘tripped the beam’), and the race officially started. The average SL was determined by dividing the length of the race by the average number of strides taken during the race. The SR was computed by dividing the number of strides by the race time in seconds. Further, the average speed and standard error of each type of race was calculated using the distance of each race and then divided by time for each winning horse.

Data were analyzed with SAS (version 9.4, Cary, NC, USA). Using the average number of strides of each winning horse, the PROC Mixed Procedure was used to evaluate differences in SR and SL between QH and TB, along with differences between distances within each breed. Differences were considered significant at P≤0.05.

## Results

Overall, QH averaged a half of stride more per second than did TB (2.88 vs 2.34 + 0.03 strides/sec; *P*<0.001). Furthermore, SR decreased (P<0.001) as race distance increased, regardless of the breed, with the highest SR being 2.96 + 0.04 strides/s during the 101-m QH race and the lowest SR (2.23 + 0.04 strides/sec) in the 2,011.7-m Breeders’ Cup Classic (Table 1).

**TABLE 1.** Average stride rate (strides/sec) for Quarter Horses and Thoroughbreds.

|  | Breed |  |  |  | SEM |
| --- | --- | --- | --- | --- | --- |
|  | Quarter Horse (n = 10) |  | Thoroughbred (n = 10) |  |  |
| Stride rate | 2.88<sup>a</sup> |  | 2.34<sup>b</sup> |  | 0.03 |
|  | Race length (meters) |  |  |  |  |
|  | 100.6 (n = 5) | 402.3 (n = 5) | 1,207.0 (n = 5) | 2,011.7 (n = 5) |  |
| Stride rate | 2.96<sup>a</sup> | 2.81<sup>b</sup> | 2.45<sup>c</sup> | 2.23<sup>d</sup> | 0.04 |

<sup>abcd</sup>Values within a row not sharing the same superscript differ ( $P < 0.001$ ). SEM corresponds to the standard error of the mean.

Average values for SL are presented in Table 2. A greater SL was observed in TB compared to QH at all distances (P<0.001). Further, QH had the shortest SL in the 100.6-m race, with it increasing in the 402.3-m race. TB stride length did not differ significantly between distances.

**TABLE 2.** Average stride length (meters) for Quarter Horses and Thoroughbreds.

|  | Breed |  |  |  | SEM |
| --- | --- | --- | --- | --- | --- |
|  | Quarter Horse (n = 10) |  | Thoroughbred (n = 10) |  |  |
| Stride length | 5.9<sup>a</sup> |  | 7.4<sup>b</sup> |  | 0.11 |
| Race length (meters) |  |  |  |  |  |
|  | 100.6 (n = 5) | 402.3 (n = 5) | 1,207.0 (n = 5) | 2,011.7 (n = 5) |  |
| Stride length | 5.0<sup>a</sup> | 6.8<sup>b</sup> | 7.3<sup>c</sup> | 7.5<sup>c</sup> | 0.04 |

<sup>abc</sup>Values within a row not sharing the same superscript statistically differ ( $P < 0.001$ ). SEM corresponds to the standard error of the mean.

The average speeds for each breed and each race are presented in Table 3. The average speed for QH and TB did not differ when calculating the speed during the official races (which involves a standing start for QH and a running start with TB), although the average speed for each distance varied with QH racing the classic distance having the fastest average speed (P<0.001). On the other hand, TB showed a significantly faster speed in sprint versus classic distances (P<0.001).

**TABLE 3.** Average speed (m/s), time and SEM for Quarter Horses (QH) and Thoroughbreds (TB) calculated from when the race officially started for each breed; from a standing start when the starting gates open for QH and from a running start when TB completed the “run-up”.

|  | Breed |  |  |  | SEM |
| --- | --- | --- | --- | --- | --- |
|  | Quarter Horse (n = 10) |  | Thoroughbred (n = 10) |  |  |
| Speed | 16.87 |  | 17.11 |  | 0.03 |
| Race length (meters) |  |  |  |  |  |
|  | 100.6 (n = 5) | 402.3 (n = 5) | 1,207.0 (n = 5) | 2,011.7 (n = 5) |  |
| Speed | 14.76<sup>a</sup> | 18.99<sup>b</sup> | 17.68<sup>c</sup> | 16.54<sup>d</sup> | 0.04 |
| Time | 6.82<sup>a</sup> | 21.19<sup>b</sup> | 68.28<sup>c</sup> | 121.66<sup>d</sup> | 0.20 |

<sup>abcd</sup>Values within a row not sharing the same superscript statistically differ ( $P<0.001$ ). SEM corresponds to the standard error of the mean.

## Discussion

Results from the study support the hypothesis that the average SR was greatest in the racing QH and SR decreases as race distance increases (Table 1). In terms of SL, the study data suggest QH have a shorter SL than TB. In addition, as hypothesized, the shortest races with the highest SR have the shortest average SL. Similarly, as also hypothesized, as the distance of the race increased, the SL also increased, but only up to the point at which the TB race distances were reached as no differences were seen between TB racing at the sprint or classic distances (Table 2).

Though it is recognized that QH are faster than TB [8, 14], the results obtained from the calculations used in this study to determine average speed (distance divided by time) would appear to lack support for this. Despite differences in speed at every race distance, the overall average speed by both breeds were not different – thereby providing temptation to conclude that QH and TB race at similar speeds. Likewise, if one compares the world speed records at the same distance (402 m), they appear somewhat similar. Currently, the QH record is held by ‘First Moonflash’, with 20.27 seconds [15], and the TB record is held by ‘Winning Brew’, with 20.57 seconds [16]. In reality, while accurate, calculating the speed using the time of the race and distance fails to take into consideration that QH races are timed from when the starting gates begin to open and the horse is standing still while TB races are timed after horses have already started running [7] and have traversed the “run-up” which is highly variable [17]. While TB tend to be relatively constant in their speed throughtout a race and the peak speed reached often is somewhat similar to their average speed, this is not true for QH. The average speed for a QH takes into account the period in which they are standing still and have not yet begun running. Pratt [8] has estimated it can be 0.6 sec before a QH has taken a step away from the starting gates. Using that estimate, roughly 9% of the race time for the QH sprint races is spent on the very first step away from the starting gate. While constituting only 3% of the race time for the QH classic races, it still represents a period during the race when the speed of the horse is at or near 0 m/s. Even after that, QH are accelerating during the initial portion of the race whereas TB had that period of acceleration during the “run-up” before the official timing of the race began. Although QH racing at the classic distance demonstrated the fasted speed, higher peak speeds could have been achieved during the sprint races. However, peak speed measurements were not within the scope of this study.

While the general public tends to believe that a long stride is correlated strongly with greater speed, this study indicates that performance is dependent on both SL and SR. That stated, a difference in stride rate within breeds at different distances is noteworthy. At the classic distances for each breed, the average QH speed over the entire race was 2.5 m/s faster than the TB speed (even with the QH timed from a standing start). With QH racing at higher speeds than TB, it is clear that the SR plays a greater role in reaching peak speeds in short distances. This findings go in agreement with the study by Hay [18] that stated, at maximum speeds, the SL remained constant or decreased slightly, contrary to the SR that was increased with the increase in speed. In fact, the fastest horse analyzed showed the highest SR but the lowest SL, showing the important role of both factors and not only SL. Moreover, the findings of another study, showed that both variables (SR and SL) increased linearly, although SL showed a tendency to decrease [11]. Of note, with the average SR being 2.96 + 0.04 strides/s during the sprint QH races, some of the horses had over 3 strides per second – a truly amazing physical feat.

In the case of human athletes, they reach maximum speeds and performance using different strategies, Elite human sprinters have been shown to possess individual preference, whether it is a higher SR or SL [3]. Yet, Hunter *et al*. [19] concluded that SR may be the most significant factor influencing speed in short distances. They examined 28 human athletes performing repeated sprint trials. Results indicate the best individual results were obtained when they had a higher SR. In fact, a negative correlation was found between SR and SL (r = −0.70). Acknowledging that there are distance differences between and within the breeds, the differences in SR and SL average between breeds can be an adjustment in racing strategy according to race distance. In other words, if trained and raced like a QH, a TB running for distances of 402 m or less may have a similar recorded time as a QH racing the same distance. For example, QH are often not ridden every day, usually being galloped only a few days per week at most [7]. This is in contrast with the TB training in which it is common practice to gallop the horses on most days and for longer distances during each ride, developing more endurance ability required for longer races in comparison with the typical distances in QH [9]. Interestingly, in the study of Ferrari *et al*. [20], the SR was increased after six months of training mature TB.

The difference in muscle mass between QH and TB is also a likely factor in the speed difference. It is reported that more than 75% of fiber muscles are Type-II in TB [21]. However, QH have a greater proportion of muscle fibers II-X when compared to TB which, in combination with their increased muscle mass compared to TB, may explain their increased speed [7, 22, 23]. Moreover, having this larger II-X fiber proportion in QH could provide a higher muscle glycolytic power and a higher maximum speed when raced short distances. While differences in muscle fiber composition are likley to be heavily influenced by genetics, differences in training techniques may also have an influence.

The current study had some limitations. In the QH races, the distance horses ran was likely very similar to the official distance of the race as QH typically run in a straight line. In contrast, the TB races were run on an oval. Often winning horses traveled greater distances than the official race distance unless they happened to be on the rail for the entire race, thus decreasing the precision in the calculation of the average SL. In truth, the average TB SL is likely slightly greater than what is reported in this study due to that variation. This does not negate the breed differences in SL, but it is acknowledged that the difference between breeds could be even greater.

Another limitation was the difficulty of counting the short steps of QH at the beginning of the races. As mentioned previously, QH races begin as the starting gates open [14] and acceleration is dramatic. There is a short period in a QH race when the horse is standing still and has not left the gate. During that rapid acceleration, QH take several short strides [8]. However, by having 15 views of each race, the challenge associated with counting those first few strides should have been ameliorated. By contrast, TB races begin when the first horse crosses in front of the flagman or electronic beam – a short distance in front of the starting gates [7]. While the TB races officially start while the horses are already running and, hence, are taking full strides (an advantage to being able to count strides), there is some degree of uncertainty in terms of being to determine exactly when the horse crossed that line. With repeated viewing of each race by five individuals, this uncertainty was likely minimized and a meter or so difference in starting point would have only a minor impact on the number of strides taken over the longer distance of the TB races. This difference in the start and how it is timed (running versus standing start) makes comparing SR and SL during different segments of the race challenging, especially at the start of the race. It was determined that only average differences in SR and SL over the entire race could be performed accurately, as opposed to, for instance, comparing SR and SL in the last 100 m of the race (which would represent the entire race for the QH sprint races). As a result of these differences in the type of racing, comparing TB and QH over the same distance could lead to inaccurate conclusions. Therefore, it is also acknowledged that differences between breeds are confounded with distance.

Besides the novelty of reporting the amazingly high SR seen in racing QH (especially at the shorter distance), this study illustrates other considerations as it relates to other physiological systems within the racing QH. First, the study raises some potential questions regarding how an increasing SR may affect the respiratory system. A locomotor-respiratory coupling system has been described in horses cantering and galloping [24, 25]. If this coupling system remains true at high speeds, the average respiratory rate may reach 134 to 147 breaths/min in the TB and 169 to 178 breaths/min in the QH with some QH individuals in the sprint races likely taking over 3 breaths/s. This respiratory rate is between 14-20% higher in the QH than the 148 breaths/min previously reported for TB racehorses [25, 26]. Although clarifying the deeper mechanisms and effects on respiratory system are beyond the scope of this study, further studies are needed to determine the potential impact on this system due to the high SR, especially for QH.

Beyond the respiratory system, the average SR findings also have possible implications for the dynamics of the equine lower limb and hoof. During each stride, the hoof momentarily comes to a halt during the stance phase of the stride (other than minor rotations or sliding of the hoof on the ground). For racing QH taking three strides per second, this suggests that three times during each second the hoof experiences rapid deceleration as the hoof comes to a stop and then experiences rapid acceleration as the hoof leaves the ground during the swing phase. For horses previously reported to reach speeds of around 89 km/h [8, 14], it suggests that the hoof on these QH may have to reach double that speed or greater (nearly 180 km/h) at some point in the swing phase. While not the point of this project, with enhanced technology being developed, it would be interesting to determine peak speeds the hooves of racing QH achieve and determine the forces associated with such rapid acceleration and deceleration.

In conclusion, despite some limitations in methodology, differences between breeds and within breeds support that a higher average SR contributes to the higher speeds previously reported for QH. Therefore, the analysis of an equine athlete must consider both SR and SL as determinants of potential performance in speed competitions. Future work could explore how the increased respiration rates affect the integrity of the respiratory system in animals with high SR, especially in the short QH races.

## Author Contributions

Conceptualization: Brian D. Nielsen, Cara I. Robison.

Data curation: Cara I. Robison, Fernando B. Vergara-Hernandez

Formal analysis: Cara I. Robison, Fernando B. Vergara-Hernandez

Funding acquisition: Brian D. Nielsen, Aimee C. Colbath.

Investigation: Fernando B. Vergara-Hernandez, Brian D. Nielsen, Cara I. Robison, Taylor A. Fabus, Jasmin L. Kompare, Rebecca A. LeCompte Lasic.

Methodology: Brian D. Nielsen, Cara I. Robison.

Project administration: Brian D. Nielsen, Aimee C. Colbath.

Resources: Brian D. Nielsen, Aimee C. Colbath

Software: Cara I. Robison, Fernando B. Vergara-Hernandez.

Supervision: Brian D. Nielsen, Aimee C. Colbath.

Validation: Brian D. Nielsen, Aimee C. Colbath, Cara I. Robison.

Visualization: Fernando B. Vergara-Hernandez, Brian D. Nielsen, Aimee C. Colbath.

Writing – original draft: Fernando B. Vergara-Hernandez, Brian D. Nielsen

Writing – review & editing: Fernando B. Vergara-Hernandez, Brian D. Nielsen, Aimee C. Colbath, Cara I. Robison, Taylor A. Fabus, Jasmin L. Kompare, Rebecca A. LeCompte Lasic.

